# StayRose: a photostable StayGold derivative red-shifted by genetic code expansion

**DOI:** 10.1101/2024.12.13.628370

**Authors:** Will Scott, Esther Ivorra-Molla, Dipayan Akhuli, Teresa Massam-Wu, Jonathan Cook, Lijiang Song, Masanori Mishima, Allister Crow, Mohan K. Balasubramanian

**Author notes:** **Corresponding Authors** Mohan K. Balasubramanian - *Centre for Mechanochemical Cell Biology and Division of Biomedical Sciences, Warwick Medical School, University of Warwick, Coventry, UK*;, Allister Crow - *School of Life Sciences, University of Warwick, Coventry, UK*;, Masanori Mishima - *Centre for Mechanochemical Cell Biology and Division of Biomedical Sciences, Warwick Medical School, University of Warwick, Coventry, UK. Authors contributed equally (Will Scott, Esther Ivorra-Molla).

## Abstract

Photobleaching of fluorescent proteins often limits the acquisition of high-quality images in microscopy. StayGold, a novel dimeric green fluorescent protein recently monomerised through sequence engineering, addresses this challenge with its high photostability. There is now focus on producing different colour StayGold derivatives to facilitate concurrent tagging of multiple targets. The unnatural amino acid 3-aminotyrosine has previously been shown to red-shift super folder GFP upon incorporation into its chromophore via genetic code expansion. Here we apply the same strategy to red-shift StayGold through substitution of Tyrosine-58 with 3-aminotyrosine. The resultant red fluorescent protein, StayRose, shows 530 nm excitation and 588 nm emission peaks, shifting from the 497 nm and 504 nm excitation and emission peaks of StayGold. StayRose also retains the favourable photostability of StayGold and can be similarly monomerised using mutations at the dimer interface. A high-resolution crystal structure of StayRose confirms the modified structure of the amino-chromophore within an unperturbed 3D fold. Although reliant on genetic code expansion, StayRose provides an important step towards developing red-shifted StayGold derivatives.

## TEXT

StayGold is a green fluorescent protein with extremely high photostability. Engineered from a *Cytaeis uchidae* protein first isolated in 2022, its minimal photobleaching under prolonged excitation makes StayGold valuable in time-lapse microscopy.^1-5^ Although the structural basis for its photostability remains unknown, we previously reported the crystal structure of StayGold,^2^ an 11-strand β-barrel with a chromophore covalently formed by G57-Y58-G59 residues in a central α-helix, alongside a buried chloride ion that interacts with several amino acids within the β-barrel. Different colour photostable StayGold derivatives would allow multi-target imaging over long timeframes. Many red fluorescent proteins have one extra double-bond than green proteins at the N-Cα bond of the first chromophore residue, which extends the π-conjugation system of delocalised electrons to cause a red-shift.^6^ Replacing tyrosine at the sfGFP chromophore (S65-Y66-G67) with an unnatural amino acid, 3-aminotyrosine (Fig. 1A), shifts fluorescence into the red range, due to a lone electron pair introduced by the amino group.^7-9^ Unnatural amino acids can be incorporated into proteins via genetic code expansion, which employs orthogonal tRNA synthetases and tRNA to recode the amber stop codon (UAG) for the chosen amino acid.^10^ We hypothesised that incorporating 3-aminotyrosine into the StayGold chromophore might produce a similar red-shift without compromising high photostability.

**Figure 1.**
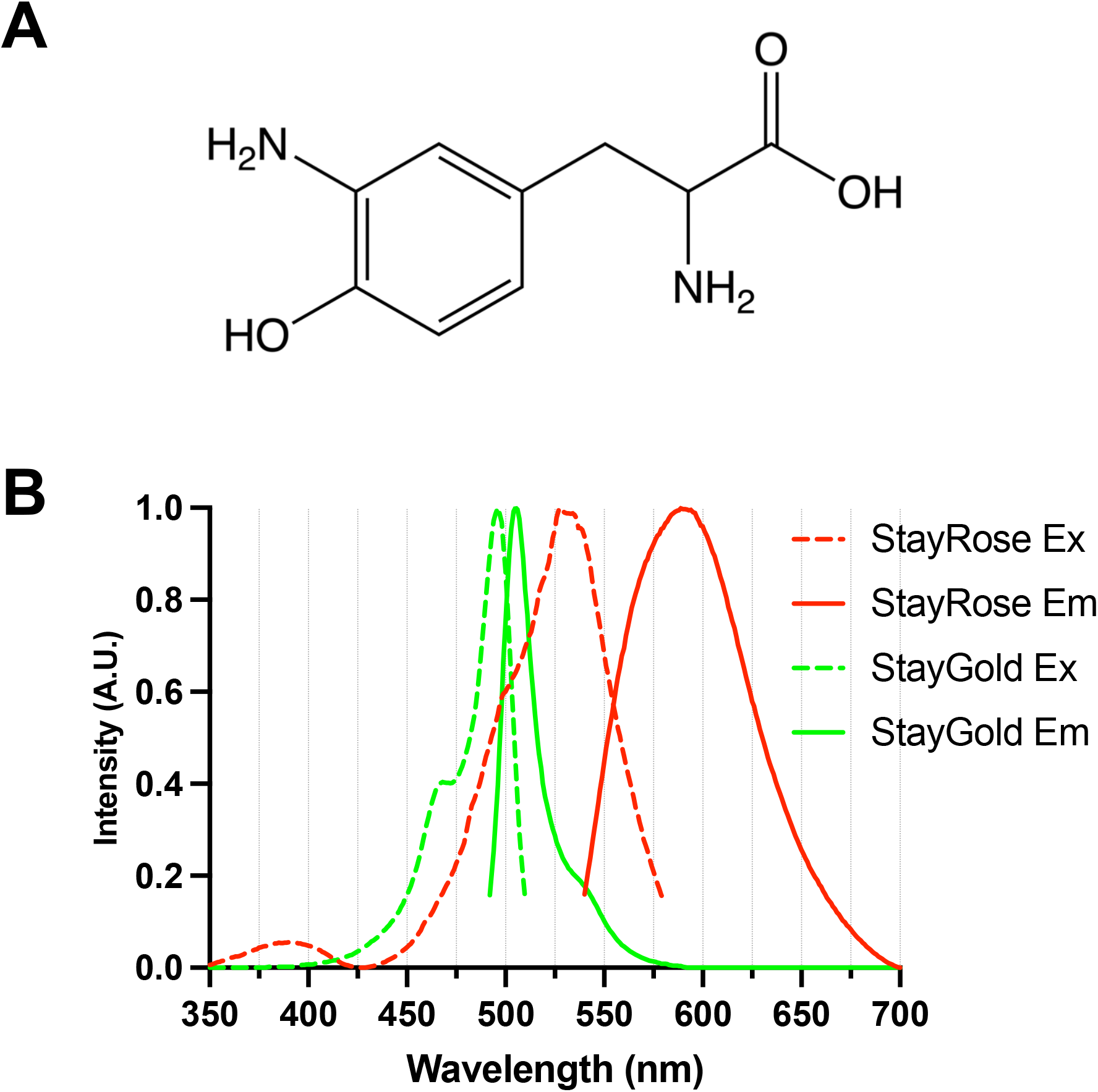
StayRose is StayGold red-shifted via 3-aminotyrosine incorporation at Y58. (A) Structure of 3-aminotyrosine. (B) Comparison of StayRose and StayGold excitation and emission fluorescence spectra. StayRose has an excitation peak of 530 nm and an emission peak of 588 nm, while StayGold has an excitation peak of 497 nm and an emission peak of 504 nm. StayGold spectra are from FPbase.^1^

3-aminotyrosine incorporation was trialled at Y58 in StayGold, which is equivalent to Y66 in sfGFP, resulting in visibly red bacterial cultures. His-tagged StayGold bearing 3-aminotyrosine at Y58, which we named StayRose, was purified (Fig. S1) and 3-aminotyrosine incorporation was confirmed via mass spectrometry of tryptic peptide fragments (Fig. S2A). StayGold exhibits peak absorbance at 497 nm and emits fluorescence with a peak at 504 nm.^2^ StayRose shows a peak absorbance at 530 nm, accompanied by a small shoulder around 497 nm (Fig. 1B), indicating the presence of a minor StayGold fraction from promiscuous incorporation of tyrosine. When excited at 530 nm, StayRose emits red fluorescence with a peak at 588 nm. The intensity of the 497 nm absorbance peak varied between preparations and in one instance was comparable to the 530 nm peak (Fig. S3A). Excitation of this particular StayRose preparation with blue light (488 nm) results in fluorescence peaking at 504 nm, with a tail extending beyond 600 nm (Fig. S3B).

The variable fraction of StayGold species within StayRose preparations was further confirmed by mass spectrometry (Fig. S2B). A minor peak 15 Da smaller than the main peak corresponds to the tyrosine-derived chromophore, with a lower mass than the 3-aminotyrosine-derived variant. Another peak, 20 Da larger than the main peak, corresponds to the nascent protein before chromophore formation via dehydration (-18 Da) and dehydrogenation (-2 Da). Sample aeration for 84 h removed the nascent peak, hypothesised to be due to completion of chromophore maturation via facilitation of oxidation. The size of the nascent peak relative to the main peak also showed preparation-dependent variation. Consistent with this, the peak absorbance/extinction coefficient of StayRose samples, based on the total protein concentration, varied from 2.5 to 3.5 x 10^4^ M^-1^ cm^-1^. The quantum yield (QY) of StayRose fluorescence, excited at 530 nm, was determined using mCherry (QY = 0.22)^11^ as the standard. The measured StayRose QY values (0.13-0.34) varied between preparations, but compare favourably to the QY reported for 3-aminotyrosine-bearing sfGFP (0.037).^9^

To further characterise StayRose, we determined its crystal structure at high resolution (Fig. 2, Table S1). StayRose crystals are intensely red, but otherwise have the same morphology as StayGold crystals (Fig. 2A) and the overall structure of StayRose is virtually identical to StayGold (Fig. 2B). An omit map showing unbiased density for the amino-modified chromophore confirms that the amino group is well defined, of high occupancy, and locked in a single conformation. The chromophore makes hydrogen bonds with Y64, E211 and N137 residues, but the amino group itself does not form any hydrogen bonds (Fig. 2C). The StayRose structure confirms that 3-aminotyrosine is comfortably accommodated by the StayGold fold and that the red-shift is due solely to the additional amino group with no further modifications elsewhere in the chromophore. These are the first structural observations of any 3-aminotyrosine-based chromophore.

**Figure 2.**
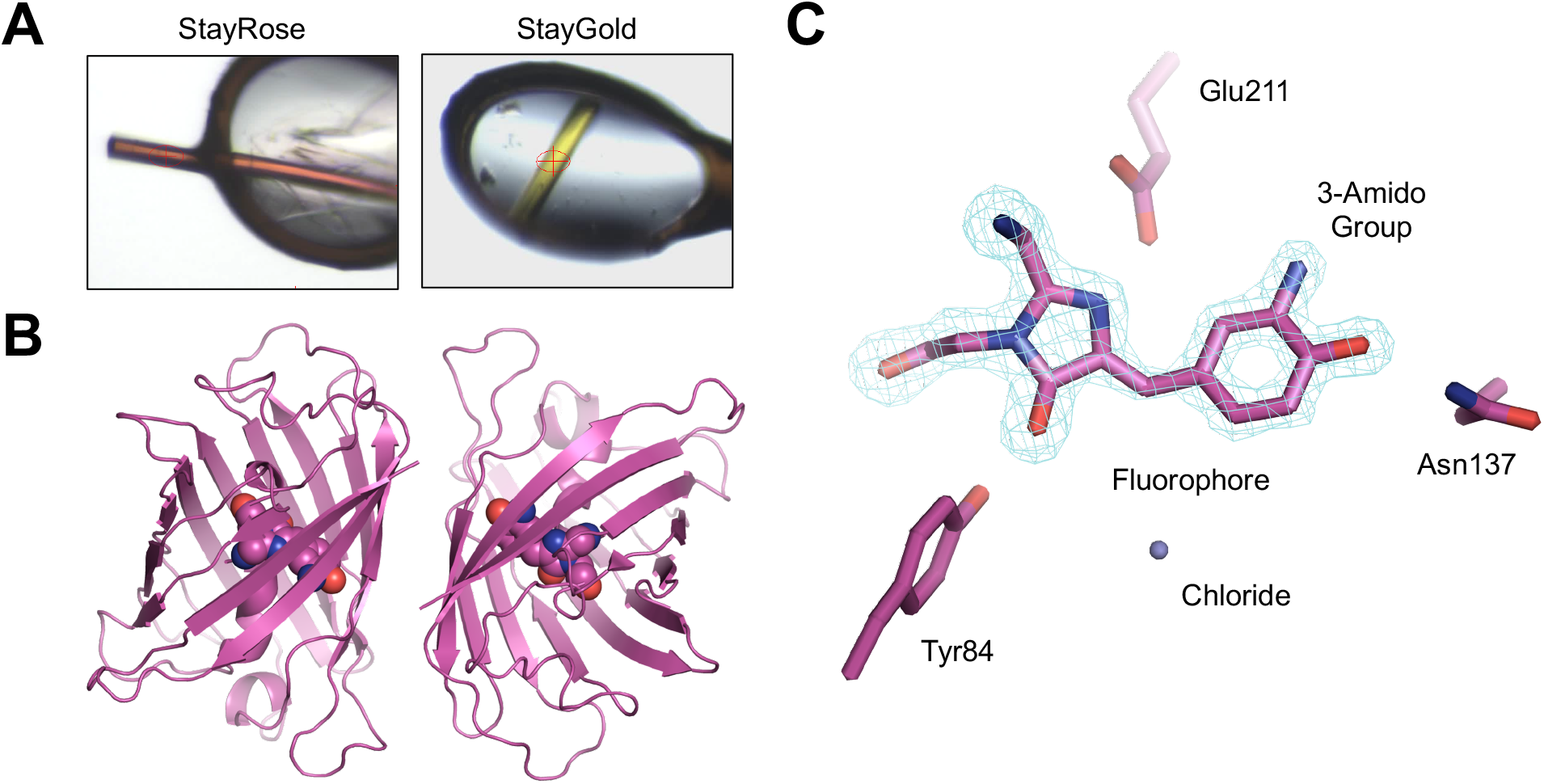
1.6 Å crystal structure of StayRose. (A) Crystals of StayRose and StayGold frozen in loops prior to X-ray data collection. (B) Structure of the StayRose dimer. The chromophore is shown as atomic spheres. (C) Close-up view of the amino-modified chromophore. A 5σ-contoured omit map shows quality of underpinning electron density.

StayGold is an obligate dimer, which can affect fusion localisation and function. Three studies have reported different monomerising sequence alterations.^2,12-13^ We reported a single residue change, E138D, that achieved monomerisation and maintained photostability.^2^ StayRose was mutagenised to produce mStayRose(E138D), which was then purified (Fig. S1). The mStayRose(E138D) variant was confirmed to be monomeric by size exclusion chromatography, while StayRose showed dimerisation (Fig. 3). Interestingly, the emission spectra of mStayRose(E138D), which showed absorbance peaks at 497 nm and 530 nm comparable to StayRose (Fig. S3C), did not exhibit an extended red fluorescence tail as was observed for dimeric StayRose under 488 nm excitation (Fig. S3B), but otherwise retained similar absorption and emission properties. This suggests that the red tail is likely due to Förster resonance energy transfer between a minority fraction of StayGold-StayRose heterodimers present within the StayRose sample, which can be resolved by monomerisation.

**Figure 3.**
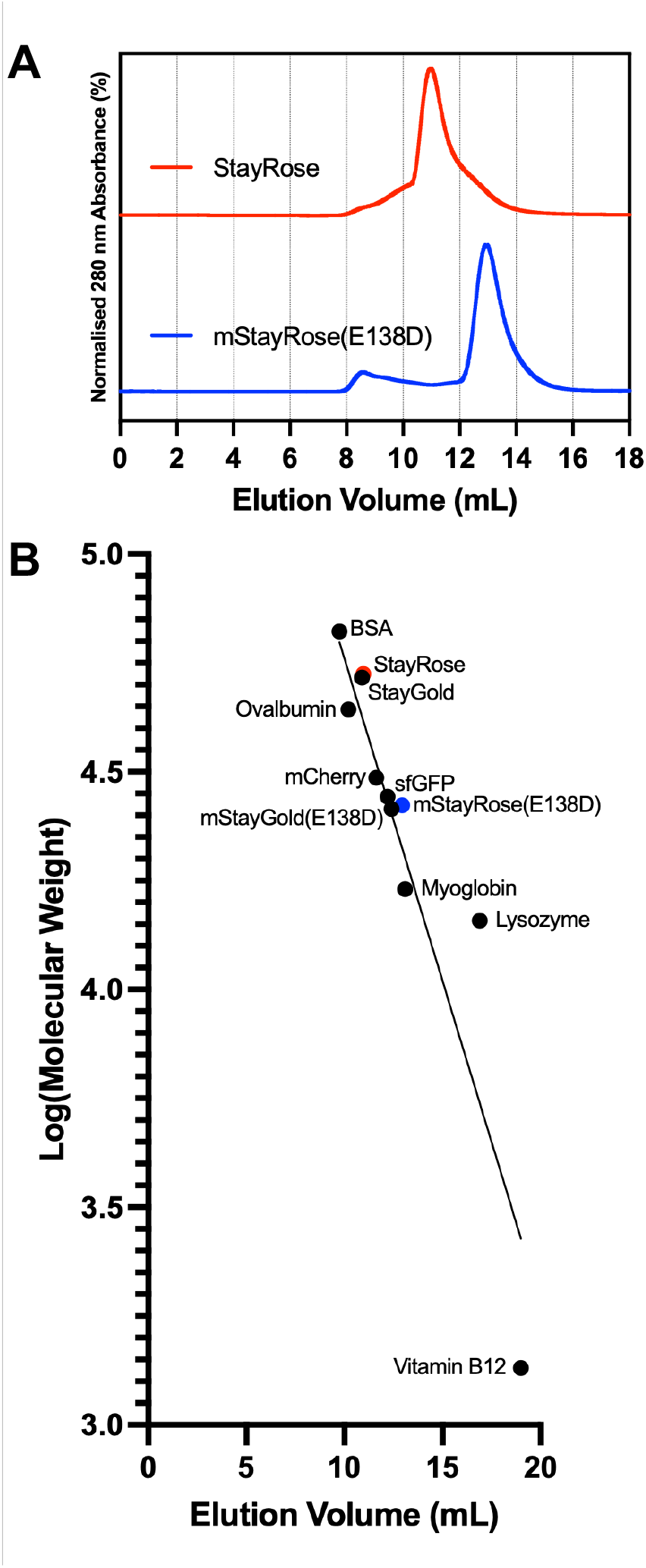
mStayRose(E138D) is monomeric and StayRose is dimeric. (A) StayRose elutes earlier than mStayRose(E138D) during size exclusion chromatography, suggesting a larger molecular size. (B) Comparison to size exclusion chromatography standards gives mStayRose(E138D) and StayRose molecular weights of 26 kDa and 52 kDa respectively, implying the former is monomeric and the latter is a dimer.

*In vitro* photostability assays of gel-immobilised StayRose and mStayRose(E138D) proteins showed both to be photostable with only modest reductions in fluorescence intensity observed under laser illumination (Fig. 4A-B). StayRose fluorescence was unexpectedly observed to increase during the first minute of laser exposure. This could be related to oxidation and protein maturation induced by laser exposure, or represent an unexpected photoactivation property of the 3-aminotyrosine-based chromophore. Purified mCherry (Fig. S1) rapidly lost brightness in the first 100 seconds, dropping 80% in intensity. As expected, the peak fluorescence intensity of mCherry was brighter than StayRose, influenced by a laser and filter set optimised for mCherry. A lower fluorescence intensity was observed for mStayRose(E138D) than StayRose. The E138D mutation is known to slightly lower the QY of StayGold from 0.92 to 0.87,^2^ but of the three reported StayGold monomers, mStayGold(E138D) has the highest QY and the second highest molecular brightness.^14^ *In vivo* bacterial photostability assays provide further evidence for the photostability of StayRose and mStayRose(E138D) (Fig. 4C-D), although the expression level of mStayRose(E138D) was far lower than StayRose and mCherry. StayRose shows a gradual increase in fluorescence, mStayRose(E138D) mostly maintains its starting intensity, and mCherry quickly photobleaches. Differences between *in vivo* and *in vitro* photostability likely reflect expression variability in bacteria, as well as the two different microscopy setups: constant spot illumination in TIRFm (*in vitro* assays), and intermittent spot illumination at in spinning disk microscopy (*in vivo* assays).

**Figure 4.**
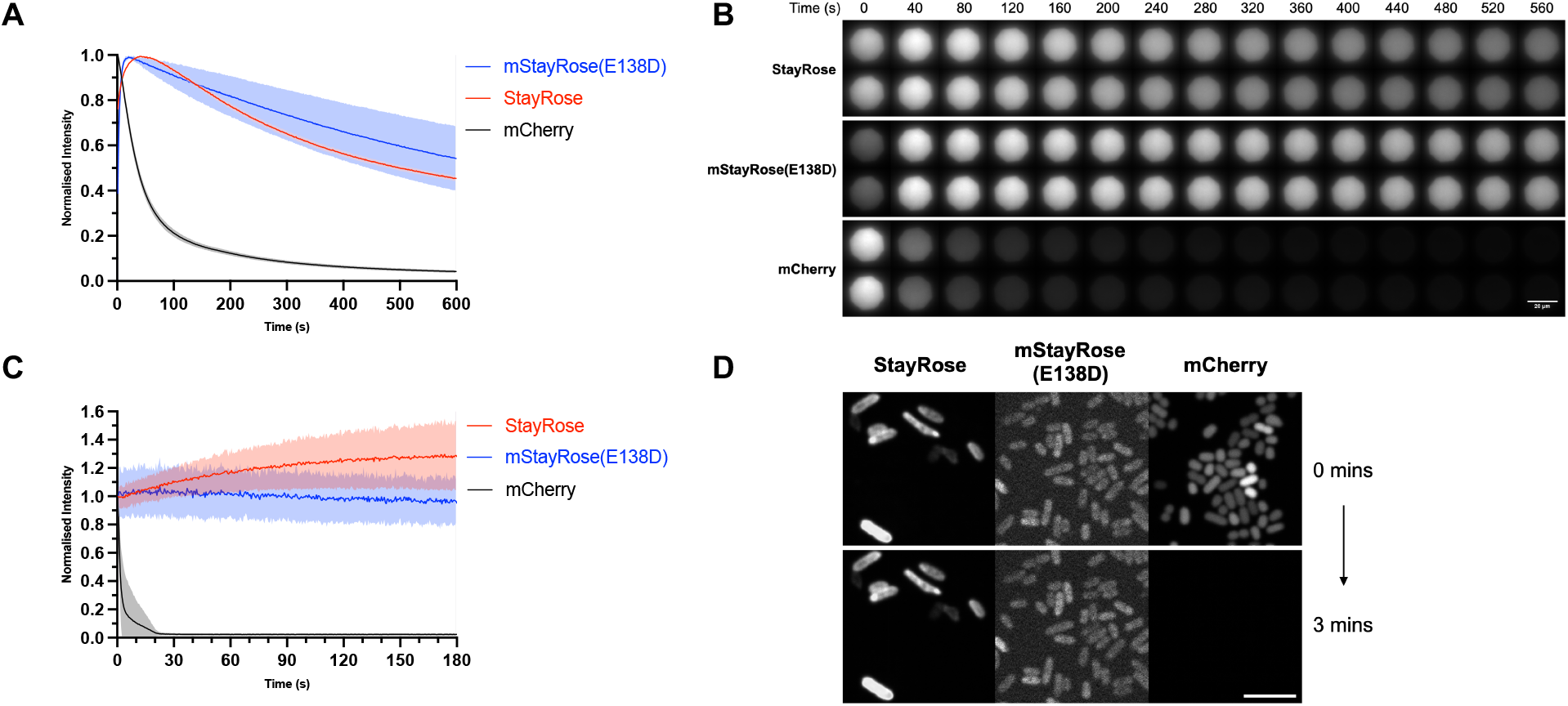
StayRose and mStayRose(E138D) are highly photostable. (A) *In vitro* photostability assays show that StayRose and mStayRose are photostable, and gain intensity during initial periods of laser exposure, while mCherry quickly loses intensity. mStayRose(E138D) had repetition variability. Intensity measurements are normalised against peak values for each protein (n = 12). (B) Representative time-lapse images of fluorescent proteins during *in vitro* assays. Intensity is normalised by peak intensity and the scale bar is 20 μm. (C) *In vivo* bacterial photostability assays showed that StayRose and mStayRose(E138D) are photostable, while mCherry photobleaches quickly. Intensity measurements are normalised to starting intensity (n = 100 cells). (D) Representative images of *in vivo* assays. Intensity is normalised by peak fluorescence and the scale bar is 5 μm. All error bars show standard deviation.

In summary, we have modified the StayGold chromophore with 3-aminotyrosine to create a new fluorescent protein, StayRose, with significantly red-shifted excitation and emission spectra. We obtained a high-resolution X-ray structure of StayRose, showed that it retains the high photostability of StayGold, and monomerised it using the E138D mutation. Several additional steps are needed to bring StayRose to the point it can be used as an effective fluorescence imaging tool. mStayRose(E138D) is a less perturbing tag than StayRose, but needs improved brightness and bacterial expression levels. StayGold itself is known to have low bacterial expression levels, and there is ongoing research across the field to improve StayGold expression and brightness, including addition of flanking terminal sequences and further amino acid substitutions. We added short tails to both ends of mStayRose, n1 and c4 sequences, respectively, which improved the brightness of expressing bacteria (Fig. S3D), consistent with previous findings in StayGold.^12,14^ There are known limitations associated with genetic code expansion-based tools, such as the requirement for multiple components reducing ease of use. Some studies have addressed problems using single plasmid expression constructs.^15^ The presence of low levels of StayGold due to tRNA synthetase infidelity is a problem for multi-target imaging studies. A community effort to improve 3-aminotyrosine incorporation efficiency would make StayRose a more powerful tool. It would be beneficial to further red-shift the 530 nm excitation peak of StayRose closer to widely used 561 nm lasers. Inspired by the T203Y mutation in sfYFP,^16^ we found that a K192Y mutation of mStayGold(E138D) results in a 4 nm excitation peak shift to 501 nm and an 8 nm emission peak shift to 512 nm (Fig. S3E). Although a small shift, point mutations like this could red-shift StayRose further. StayRose demonstrates that the structural features crucial for StayGold’s photostability are not limited to green fluorescence, marking an important step towards developing photostable StayGold variants in a spectrum of colours.

## ASSOCIATED CONTENT

### Supporting Information

The Supporting Information file is available free of charge.

Material and Methods, Table S1, Figures S1-S3 (PDF).

### Accession Codes

StayGold, PDB 8BXT; sfGFP, NCBI UFQ89826; StayRose, PDB 9G7Q; mCherry, NCBI UFQ89828.

## AUTHOR INFORMATION

### Author Contributions

The manuscript was written through contributions of all authors. All authors have given approval to the final version of the manuscript. ^‡^These authors contributed equally.

### Notes

Authors declare no competing financial interests.

## ACKNOWLEDGMENTS

The authors thank: staff at Diamond Synchrotron; staff at the University of Warwick Computing and Advanced Microscopy Unit (CAMDU); Rob Cross (University of Warwick) for access to fluorimetry equipment; Cleidi Zampronio and the Warwick Proteomics Research Technology Platform; Gregory Campbell, Andrew Hart and Arthur Margolis for pilot experiments during undergraduate projects.

## ABBREVIATIONS

QY: quantum yield

## Table of Contents Graphic

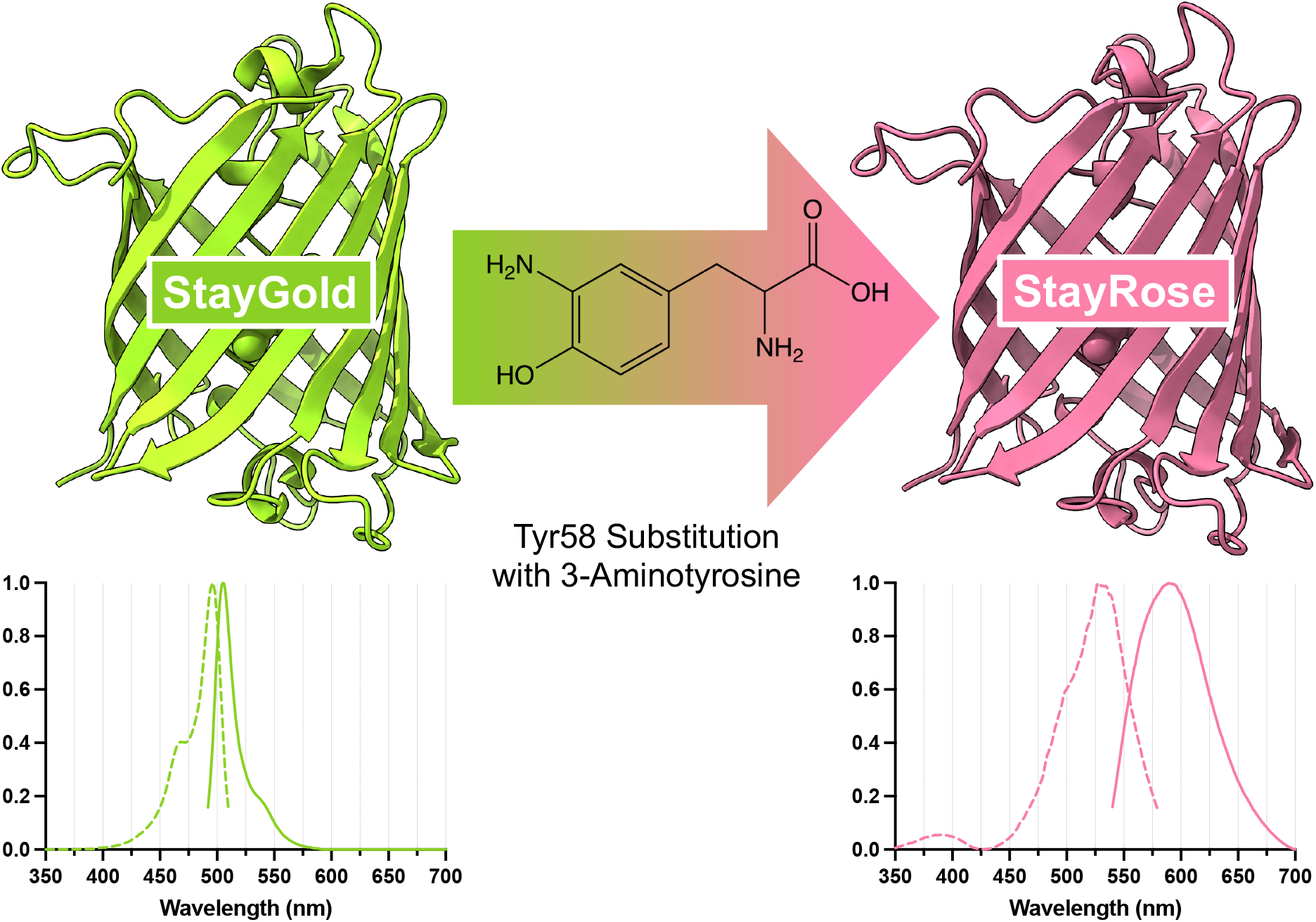

