## Supporting Information for "StayRose: a photostable StayGold derivative red-shifted by genetic code expansion"

Material and Methods
Supplemental Table
- Table S1. Crystallographic data collection and refinement statistics
Supplemental Figures
- Figure S1. His-tagged fluorescent protein purification
- Figure S2. Mass spectrometry analyses of purified StayRose and mStayRose(E138D)
- Figure S3. Fluorescence of novel StayGold derivatives

### Materials and Methods

#### Cloning, expression & purification of fluorescent proteins

pET-MCN-10His-StayRose and pET-MCN-10His-mStayRose(E138D) plasmids were generated by substituting Y58 with an amber stop codon (TAG) via mutagenesis of pET-MCN-10His-StayGold (Addgene #211360) and pET-MCN-10His-mStayGold(E138D) (Addgene #211361) respectively. pET21a-6His-mStayGold(E138D+K192Y) plasmid was generated via cloning of mStayGold(E138D) from pET-MCN-10His-StayGold into a pET21a vector, followed by K192Y mutagenesis. Also used was pBAD-6His-mCherry (AddGene #54630). For incorporation of 3-aminotyrosine in all genetic code expansion experiments, pEvol-MjaYRS (AddGene #153557) plasmid was used. All sequences were confirmed via nanopore whole plasmid sequencing.

His-StayRose, His-mStayRose(E138D), His-mStayGold(E138D+K192Y), and His-mCherry proteins were purified. For this, each of the fluorescent protein reporter constructs were transformed into BL21(DE3) *Escherichia coli*, with a further transformation of pEvol-MjaYRS plasmid carried out for those requiring 3-aminotyrosine incorporation. Liquid cultures were grown up shaking at 36°C. For cultures requiring 3-aminotyrosine incorporation, 100 mM 3-aminotyrosine (Bachem, 4027898) in 0.2 M HCl was added to a final concentration of 1 mM 3-aminotyrosine in OD 0.2 cultures. OD 0.6 liquid cultures were induced for 28 h at 18°C with 0.5% Arabinose and/or 0.9 mM IPTG as appropriate. Resulting bacterial pellets were lysed via sonication in lysis buffer (50 mM phosphate buffer, 300 mM NaCl, 0.1 mM $MgCl_2$, 10 mM imidazole, 0.3 mM PMSF and cOmplete™ Protease Inhibitor Cocktail (pH7.5)). HisPur Ni-NTA resin (Thermo Scientific, 88222) was used to purify proteins in wash buffer (50 mM phosphate buffer, 500 mM NaCl, 30 mM imidazole (pH 7.5)) before elution in elution buffer (50 mM phosphate buffer, 500 mM NaCl, 500 mM imidazole (pH7.5)). Sample buffer was exchanged in PD midiTrap G-25 desalting columns (Cytiva, 28918008) for storage buffer (20 mM HEPES and 150 mM NaCl (pH 7.5)). Purified sfGFP, StayGold, and mStayGold(E138D) protein samples used in this study were the same preparations used in Ivorra-Molla *et al.* (2023).[2]

#### Digested protein mass spectrometry analysis

Purified protein samples were run on a 12% SDS-PAGE acrylamide gel and Coomassie blue stained, from which bands were excised. Bands were de-stained in 50% ethanol and 50 mM ammonium bicarbonate (ABC) (3 x 20 minute washes at RT), dehydrated in ethanol (5 minutes at RT), and reduced in 40 mM CAA and 10 mM TCEP (5 minutes at 70°C). Following further washing in 50% ethanol and 50 mM ABC (3 x 10 minute washes at RT) and dehydration in ethanol (5 minutes at RT), gels were hydrated in 2.5 ng/µL trypsin and incubated overnight at 37°C. The liquid fraction was collected and further peptide was extracted from the gel via water bath sonication (3 x 10 minutes at RT) in 25% acetonitrile and 5% formic acid with liquid fraction collection after each round. Combined liquid fractions were concentrated via speed vacuum and resuspended in 50 µL 2% acetonitrile and 0.1% trifluoroacetic acid.

Reversed phase chromatography was used to separate tryptic peptides prior to mass spectrometric analysis. Two C18 columns were utilised, an Acclaim PepMap µ-precolumn cartridge 300 µm i.d. x 5 mm 5 µm 100 Å (Thermo Fisher Scientific) and a 75 µm x 40 cm 1.9 µm (Bruker nanoElute Forty Analytical column). The columns were installed on a NanoElute UHPLC system (Bruker Daltonics, Germany). Mobile phase buffer A was composed of 0.1% formic acid in water and mobile phase B 0.1% formic acid in acetonitrile. Samples were loaded on onto the µ-precolumn equilibrated in 2% aqueous acetonitrile containing 0.1% formic acid and peptides were eluted onto the analytical column at 350 nL min$^{-1}$ by increasing the mobile phase B concentration from 3% B to 15% over 31 min, then to 35% B over 10 min, and to 85% B over 3 min, followed by a 10 min re-equilibration at 0% B. NanoElute was coupled online to a hybrid timsTOF Pro (Bruker Daltonics, Germany) via a CaptiveSpray nano-electrospray ion source.[17] The timsTOF Pro was operated in Data-Dependent Parallel Accumulation-Serial Fragmentation (PASEF) mode. Peptides were separated by ion mobility depending on their collisional cross sections and charge states. The method settings were as follows: mass range 100 to 1700 m/z, ion mobility range $1/K_0$ Start 0.6 Vs/cm$^2$ End 1.6 Vs/cm$^2$, Ramp rate 9.42 Hz and Duty cycle 100%. Peptide datasets were searched against protein specific databases using Mascot Database Search by Matrix Science, with up to two missed cleavages, fixed modification of cysteine carbamidomethylation and variable modification of methionine oxidation, as well as UAA modification where applicable.

**Whole protein mass spectrometry analysis**
100 µg purified StayRose and mStayRose(E138D) protein samples were buffer exchanged in a 10 kDa cut-off 0.5 mL Amicon centrifugal filter device to 500 µL 35 mM ammonium acetate (pH 7.4). Mass spectrometry analysis was carried out using a Bruker MaXis II coupled with Dionex 3000RS UHPLC with C4 column (100 x 2.1 mm, 2.7 µm), a gradient of 5-100% Water/CAN, and a flow rate of 0.2 mL/minute. The 84 h matured sample was aerated for 84 h to allow protein maturation prior to mass spectrometry.

**Structure determination using x-ray crystallography**
Crystallisation screens used StayRose in 150 mM NaCl, 20 mM HEPES (pH 7.5) in MRC 2-drop plates. Sitting drops were set using a Formulatrix NT8 robot with 1 µL drops of 1:2 or 2:1 protein solution and crystallisation reagent. Crystals were obtained in the SG1 HT96 Eco Screen (Molecular Dimensions) condition H4 (25% PEG (Polyethylene glycol) 3350, 0.1 M Bis–Tris (pH 6.5) and 0.2 M sodium acetate).

StayRose crystals were cryoprotected with 30% glycerol. X-ray diffraction experiments were conducted remotely at beamline I04 at the Diamond Synchrotron. Diamond's auto-processing pipeline integrated diffraction images using Dials.[18] The CCP4 suite[19] was used to perform subsequent processing, with AceDRG[20] used to generate constraints for the 3-aminotyrosine ligand. Aimless[21] was used for scaling and space group determination. Phases were determined by molecular replacement using Phaser.[22] Model building and rebuilding used Coot[23] and refinement was performed with REFMAC.[24] The model was validated using tools from PROCHECK,[25] Rampage[26] and Coot.[23] Images were generated using PyMOL.[27] The StayRose structure was deposited in the protein databank with accession code 9G7Q.

**Fluorescence excitation and emission spectra**
A Cary Eclipse fluorescence spectrophotometer with a 5 nm excitation and emission slit width was used to acquire fluorescence spectra. Spectra of three serial (100, 50 and 25%) dilutions of each protein were taken using a 600 nm min$^{-1}$ scan rate at 1 nm intervals.

**Extinction coefficient determination**
A Cary 50 Conc UV Visible spectrophotometer was used to measure absorbance of each protein at four serial (100, 75, 50 and 25%) dilutions. The molar extinction coefficients ($\varepsilon_{FP}$) were calculated using the peak absorbance ($A_{peak}$), the absorbance at 280 nm ($A_{280}$) and the theoretical extinction coefficients at 280 nm ($\varepsilon_{280}$) (with one tyrosine omitted from the sequence, https://web.expasy.org/protparam/).[28]

$$\varepsilon_{FP} = \varepsilon_{280} \times A_{peak}/A_{280}$$

**Quantum yield measurements**

Fluorescent protein quantum yields were determined by comparing the ratio between the integrated fluorescence emission and the absorbance at the peak excitation for the fluorescent protein, with the same ratio for mCherry.[28] The reported quantum yield value for mCherry of 0.22 was used for this.[11] The fluorescence/absorbance ratios were each determined using the gradient from a plot of integrated fluorescence versus absorbance.

**Size exclusion chromatography**

An ÄKTA Pure FPLC with a Superdex 75 Increase 10/300 GL column was used to perform size exclusion chromatography, with a buffer of 20 mM HEPES and 150 mM NaCl (pH 7.5) and a 0.8 mL min$^{-1}$ flow rate. For each run, ~1 mg mL$^{-1}$ (38 µM) protein was loaded onto the column via a 0.5 mL loop. As well as the novel fluorescent proteins, additional runs of bovine serum albumin (66 kDa) and hen egg white lysozyme (14 kDa), StayGold (52 kDa (as a dimer)), mStayGold(E138D) (26 kDa), mCherry (31 kDa), sfGFP (27 kDa), and BIORAD molecular weight standards (thyroglobulin (670 kDa); ɣ-globulin (158 kDa); ovalbumin (44 kDa); myoglobin (17 kDa); and vitamin B12 (1.35 kDa)) were used for estimation of protein molecular weights.

**Microscopy and *in vivo* photostability**

Liquid cultures of BL21(DE3) *E. coli* were transformed with the fluorescent protein expression constructs described above (and pEvol-MjaYRS plasmid where appropriate), treated with 3-aminotyrosine (if required) and induced as per the steps described in the protein purification section. Induced bacteria were imaged on microscopy slides, using a spinning disk confocal microscope (Andor Revolution XD imaging system, equipped with a Nikon ECLIPSE Ti inverted microscope, a confocal unit Yokogawa CSU-X1, an Andor iXon Ultra 888 EMCCD camera, a ×100 oil immersion 1.45 NA Nikon Plan Apo Lambda objective (69 nm/pixel), and Andor IQ acquisition software). Cells were imaged for 3 minutes at 0.5s intervals with a 60% power (1.19 mW) 561 nm laser. The microscope area of illumination is 11748.7 µm$^2$, corresponding to an overall illumination of 10.1 W cm$^{-2}$. A set area of fluorescence intensity from each cell was measured from every slice using the image processing software Fiji (https://imagej.net/software/fiji/). Intensity measurements were normalised to the first data point.

***In vitro* photostability**

Photostability of purified proteins was measured using the method described in Cranfill *et al.* (2016).[28] In brief, fluorescent proteins were embedded in a polyacrylamide gel (20% acrylamide (mono:bis 29:1), 150 mM NaCl and 20 mM Tris (pH 7.5)), which was polymerised between a 1.5 coverslip and a glass slide with 15.5 µm polystyrene beads (Bangs Laboratory, PS07N) as spacers. The samples were observed using a TIRFm setup, namely a Nikon Eclipse Ti-E/B microscope equipped with a Ti-E TIRF illuminator (561 nm CW laser line), a Zyla sCMOS 4.2 camera, a ×100 oil immersion 1.49 NA Nikon CFI Apo TIRF objective lens, and Andor iQ3 software. A 381.9 µm$^2$ octagonal area within each sample was continuously illuminated using a 561 nm laser beam at 100% intensity and a 0° angle of incidence at 1s intervals for 10 minutes. The laser power was measured as 347 µW, corresponding to illumination at 90.9 W cm$^{-2}$.

**Supplementary Tables**

**Table S1. Crystallographic data collection and refinement statistics.**

| | **StayRose (9G7Q)** |
|---|---|
| **Data collection** | |
| Beam line | Diamond I04 |
| Wavelength (Å) | 0.95373 |
| **Crystal parameters** | |
| Space group | $P6_1$ |
| Unit cell dimensions (Å) | 133.31, 133.31, 58.35 |
| Unit cell angles (°) | 90, 90, 120 |
| **Reflection data*** | |
| Resolution range (Å) | 44.63-1.65 (1.68-1.65) |
| Unique reflections | 70,220 (3,349) |
| $R_{sym}$ | 0.150 (1.423) |
| $R_{pim}$ | 0.033 (0.335) |
| I/σ(I) | 14.3 (2.0) |
| $CC_{½}$ | 0.999 (0.784) |
| Completeness (%) | 98.5 (95.4) |
| Multiplicity | 20.7 (18.3) |
| Wilson B (Å$^2$) | 15.8 |
| **Refinement†** | |
| Resolution (Å) | 43.67 – 1.65 |
| Number of reflections | 66,707 |
| $R_{overall}$ | 0.155 |
| $R_{free}$ | 0.177 |
| Rms (bond lengths) (Å) | 0.011 |
| Rms (bond angles) (°) | 1.76 |
| **Model B-factors** | |
| StayGold (chain A) (Å$^2$) | 19.5 |
| StayGold (chain B) (Å$^2$) | 19.3 |
| Waters (Å$^2$) | 30.6 |
| **Ramachandran statistics‡** | |
| Favoured (%) | 98.3 |
| Allowed (%) | 1.7 |
| Outlier (%) | 0 |

Values in parentheses indicate the highest resolution bin.

Refinement statistics are from REFMAC.[24]

Ramachandran statistics as reported by Rampage.[26]

### Supplemental Figures

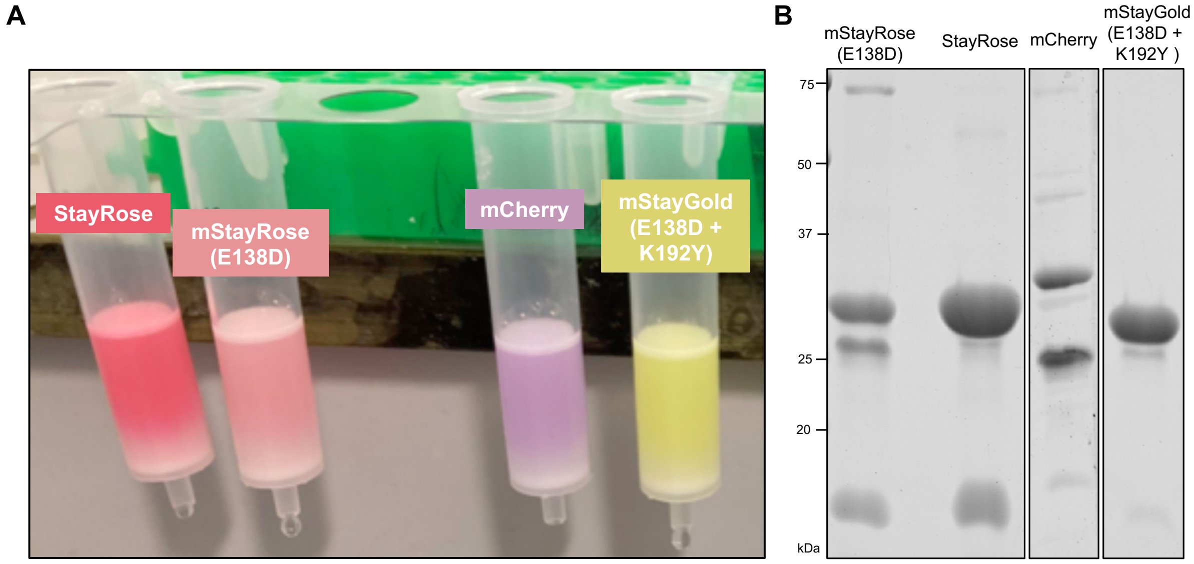


**Figure S1. His-tagged fluorescent protein purification.** (A) Image of purified fluorescent proteins loaded on PD midiTrap G-25 column during buffer exchange. Bacterial expression of mStayRose(E138D) was observed to be lower than StayRose, hence the lower intensity colour. (B) SDS-PAGE gel of purified fluorescent proteins. Genetic code expansion commonly produces two side products: a protein truncated at the amber stop codon site and a protein bearing a natural amino acid at the amber stop codon site. The former is due to a low rate of failed amber suppression and the latter due to a low level of recognition for tyrosine by the orthogonal tRNA synthetase. mCherry appears as two bands because of the double bond introduced in the main peptide chain by the matured chromophore, which breaks during sample heating prior to SDS-PAGE. This leaves immature protein at the larger full-sized band, and damaged mature protein as the lower band.[29]

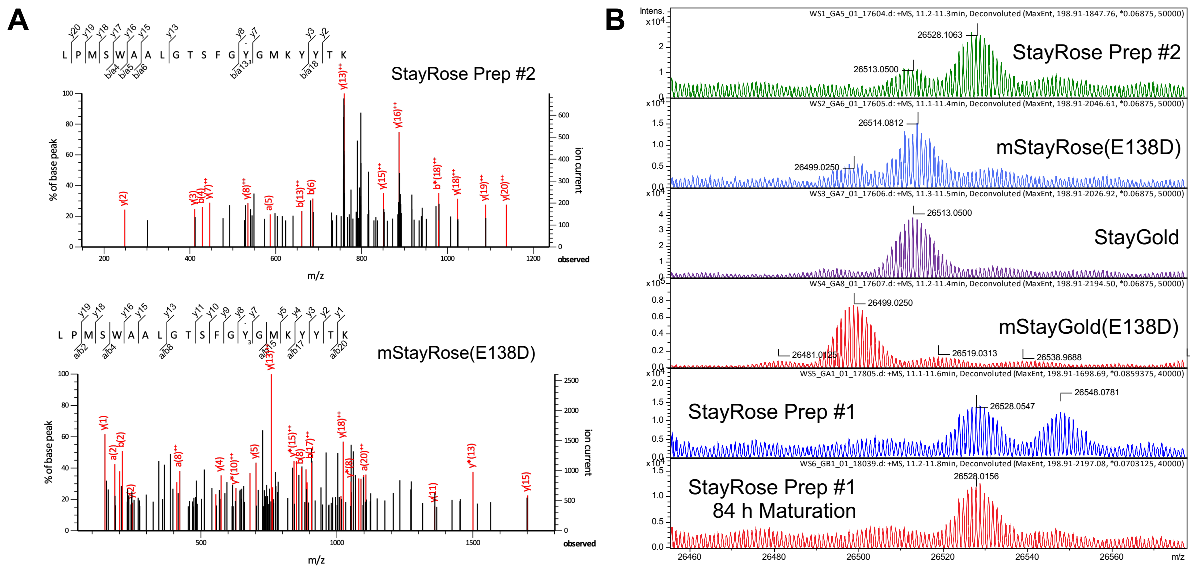


**Figure S2. Mass spectrometry analyses of purified StayRose and mStayRose(E138D).** (A) Spectra of digested StayRose and mStayRose(E138D) peptide fragments confirm 3-aminotyrosine incorporation. (B) Spectra of whole StayRose and mStayRose(E138D) proteins allow quantification of molecules incorporating 3-aminotyrosine or tyrosine at position 58, as well as comparison of nascent and matured protein peaks. The ratio of each varies between preps, indicating variability in incorporation rate.

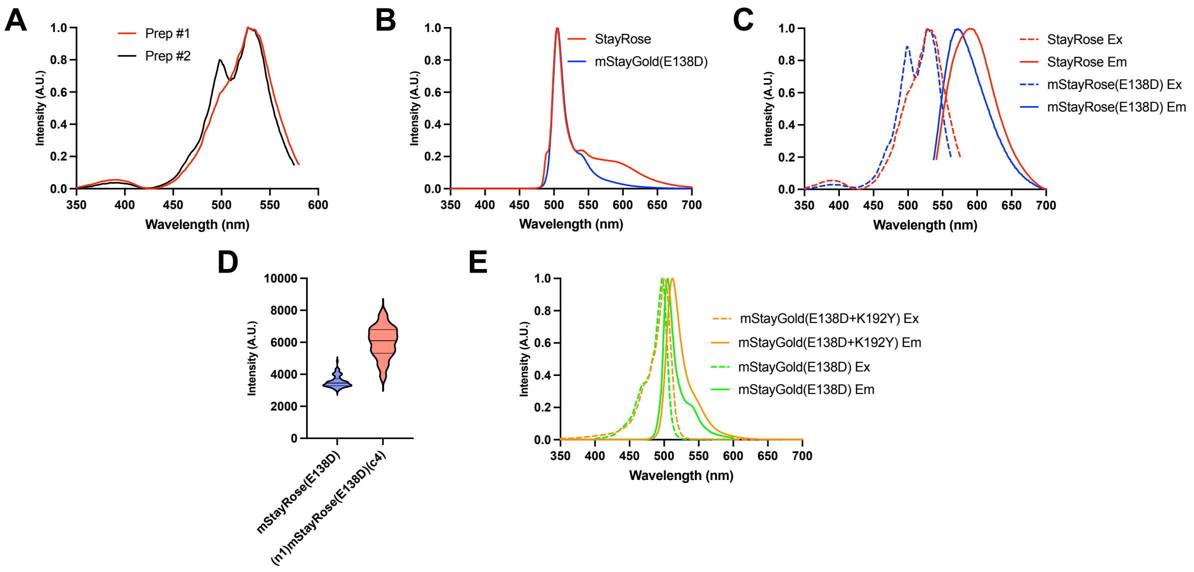


**Figure S3. Fluorescence of novel StayGold derivatives.** (A) Excitation spectra of two StayRose protein preparations show variability in 3-aminotyrosine incorporation efficiency. Emissions at 588 nm were measured. (B) Emission spectra with excitation at 488 nm of StayRose and mStayRose(E138D) show a lengthier tail beyond 600 nm for StayRose. (C) Excitation and emission spectra of StayRose and mStayRose(E138D). 588 nm emissions were detected for excitation spectra, and emission spectra were detected at 530 nm excitation. (D) (n1)mStayRose(c4)(E138D) has higher cellular brightness than mStayRose(E138D) in *E. coli* (n = 90 cells). (E) Excitation and emission spectra of mStayGold(E138D + K192Y) and mStayGold(E138D). For mStayGold(E138D + K192Y), 512 nm emissions were detected for excitation spectra, and emission spectra were detected at 501 nm excitation. mStayGold(E138D) spectra from Ivorra-Molla *et al*. (2023)[2] were used. For all spectra in this figure, n = 3.
